# Tumor-immune metaphenotypes orchestrate an evolutionary bottleneck that promotes metabolic transformation

**DOI:** 10.1101/2022.06.03.493752

**Authors:** Jeffrey West, Frederika Rentzeperis, Casey Adam, Rafael Bravo, Kimberly A. Luddy, Mark Robertson-Tessi, Alexander R. A. Anderson

## Abstract

Metabolism plays a complex role in the evolution of cancerous tumors, including inducing a multifaceted effect on the immune system to aid immune escape. Immune escape is, by definition, a collective phenomenon by requiring the presence of two cell types interacting in close proximity: tumor and immune. The microenvironmental context of these interactions is influenced by the dynamic process of blood vessel growth and remodelling, creating heterogeneous patches of well-vascularized tumor or acidic niches. We present a multiscale mathematical model that captures the phenotypic, vascular, microenvironmental, and spatial heterogeneity which shapes acid-mediated invasion and immune escape over a biologically-realistic time scale. We model immune escape mechanisms such as i) acid inactivation of immune cells, ii) competition for glucose, and iii) inhibitory immune checkpoint receptor expression (PD-L1) under anti-PD-L1 and sodium bicarbonate buffer therapies. To aid in understanding immune escape as a collective cellular phenomenon, we define immune escape in the context of six collective phenotypes (termed “meta-phenotypes”): **Self-Acidify, Mooch Acid, PD-L1 Attack, Mooch PD-L1, Proliferate Fast**, and **Starve Glucose**. Fomenting a stronger immune response leads to initial benefits but this advantage is offset by increased cell turnover that accelerates the emergence of aggressive phenotypes by inducing an evolutionary bottleneck. This model helps to untangle the key constraints on evolutionary costs and benefits of three key phenotypic axes on tumor invasion and treatment: acid-resistance, glycolysis, and PD-L1 expression. The benefits of concomitant anti-PD-L1 and buffer treatments is a promising treatment strategy to limit the adverse effects of immune escape.

Significance statement
In this work, we present a multi-scale mathematical model that captures the phenotypic, vascular, microenvironmental, and spatial heterogeneity which shapes acid-mediated invasion and immune escape over a biologically-realistic time scale. To aid in understanding immune escape as a collective cellular phenomenon, we introduce the concept of metaphenotypes: immune escape mechanisms that account for cellular phenotype and surrounding context. Metaphenotypes are defined by accounting for the following: cellular phenotype, microenvironmental factors, neighboring cell types, and immune-tumor interactions. These metaphenotypes provide insight into why targeting intratumoral pH with bicarbonate buffer is a synergistic combination treatment when paired with immune checkpoint blockade.

## 1 Introduction

Metabolism plays a complex but key role in the evolution of cancerous tumors. Localized hypoxia due to vascular instability and dysfunction leads to acidification of the tumor microenvironment. Decreased pH selects for acid-resistant tumor-cell phenotypes, followed by the emergence of aerobic glycolysis (i.e., the Warburg effect^1^). Further microenvironmental acidification by these metabolically aggressive cells foments acid-mediated invasion^2, 3^. This nonlinear evolutionary trajectory through a range of metabolic phenotypes has been studied clinically, experimentally, and theoretically^4–7^.

The effect of metabolic processes on the immune system is a multifaceted interaction between intracellular metabolism of many varied cell types with the surrounding microenvironment. Immunometabolism is a growing area of study^8^ where systems biology and mathematical approaches are highly suited to studying tumor-immune dynamics^9–15^, whether using non-spatial continuum approaches^16^ or spatial agent-based models^17^. However, very few tumor-immune models to date have incorporated the effects of cancer metabolism on immune function^18^.

### 1.1 Metabolism and the tumor-immune response

Cytotoxic T lymphocytes (CTL, also known as *α*/*β* CD8+ effector T-cells) are key players in adaptive immune response which are activated via antigen presentation during the body’s initial inflammatory response and subsequently rapidly proliferate. Programmed cell death-1 (PD-1) is an inhibitory immune checkpoint receptor expressed on activated CTLs, and programmed cell death ligand-1 (PD-L1) is a cell surface marker that activates PD-1 signaling^19^. Some cancers constitutively express PD-L1, leading to the development of anti-PD-1/PD-L1 therapy to counter this immune escape mechanism. Immune escape or evasion mechanisms may select for subclonal populations capable of withstanding immune predation^20^, often well before tumor invasion into normal tissue^21^.

We investigate two key connections between tumor metabolism and immune function: acid-inactivation and glucose competition. Acidic microenvironments have been shown to inactivate otherwise viable CTLs^22^, as cells rescued from low pH environments had the ability to regain effector function^23^. Tumor acidity also promotes regulatory T-cell (Tregs) activity as well as an increase of PD-1 expression on Tregs, indicating that PD-1 blockade may increase suppressive capacity^24^. Tumor-infiltrating CD8+ T-cells require glucose to support their killing function, hence competing for glucose with cancer cells dampens their anti-cancer response^25^. In contrast, Tregs avoid competition for glucose through rewired metabolism away from aerobic glycolysis, which enhances their immune-suppression function within the tumor^26^.

Acid-inactivation and glucose competition may diminish immunotherapy efficacy, suggesting a potential synergy between targeting intratumoral pH and immune checkpoint blockade. For example, combining oral bicarbonate buffering with immunotherapy (adoptive T-cell transfer, anti-CTLA4, or anti PD-1) increased responses in murine cancer models, presumably due to increased immune activity in a less acidic microenvironment^23^. Another study showed that targeting bicarbonate transporters (e.g. SLC4A4) known to contribute to extracellular pH during progression of pancreatic adenocarcinomas (PDAC)^27^ reduces tumor acidity, increases activation, cytotoxic activity, and perfusion of CD8+ T-cells, and sensitizes PDAC-bearing mice to immune checkpoint inhibition^27^. Mechanistic modeling has been used to investigate the treatment effects of systemic pH buffers (sodium bicarbonate) to limit microenvironmental selection for acid-adapted phenotypes arising, resulting in significantly delayed carcinogenesis in TRAMP mice^6, 28^. Buffers reduce intratumoral and peritumoral acidosis, inhibiting tumor growth^4^ and reducing spontaneous and experimental metastases^29, 30^.

### 1.2 The tumor-immune gambit

The back and forth of cancer treatment and a tumor’s evolutionary response has been compared to a chess match^31^. Similarly, we show that immune predation of tumors can be likened to an “immune gambit”, where a temporary sacrifice of (normal glycolytic) cells on the periphery leads to long-term acceleration of the invasion of aggressive (highly glycolytic) phenotypes into surrounding tissue. Vascular dynamics are often abnormal in tumors whereby areas of poor vascularization are prone to develop acidic niches. We show that poor vascularization selects for aggressive phenotypes while high vascularization undergoes low levels of evolution. This phenomena has a Goldilocks effect, which occurs only under moderate levels of immune response. The immune gambit is described as a collective phenotype, which critically depends on the interplay between local vascularization, immune infiltration, and immune evasive phenotypes (PD-L1).

### 1.3 Collective cellular phenotypes: the “Metaphenotype”

In order to describe the collective nature of phenotypes operating within the context of surrounding cells and environmental conditions, we propose the concept of a “metaphenotype”. Each of these metaphenotypes account for phenotypic traits (e.g. PD-L1 expression or glycolytic rate) as well as surrounding environmental context (e.g. local glucose or pH concentration), and competition with neighboring cell types (immune, cancer, normal). A mathematical model is the ideal testing ground for defining collective phenotypes because it enables precise characterization of local context. A simple, contrived example in figure 1 illustrates the need to quantify context-dependent selection in this model. This figure shows the time-evolution of identical phenotypic compositions that have varied initial spatial configurations (mixed or shell). The mixed configuration of low glycolysis (blue) and high glycolysis (purple) phenotypes leads to no evolution. The volumetric concentration of acid produced by aggressive cells is not enough to cause invasion when highly glycolytic cells are seeded far apart but artificially placing the aggressive high glycolysis phenotypes on the rim leads to invasion from increased volumetric acid via a group-effect. Clearly, both tumor phenotypic composition and neighboring context are important.

**Figure 1.**
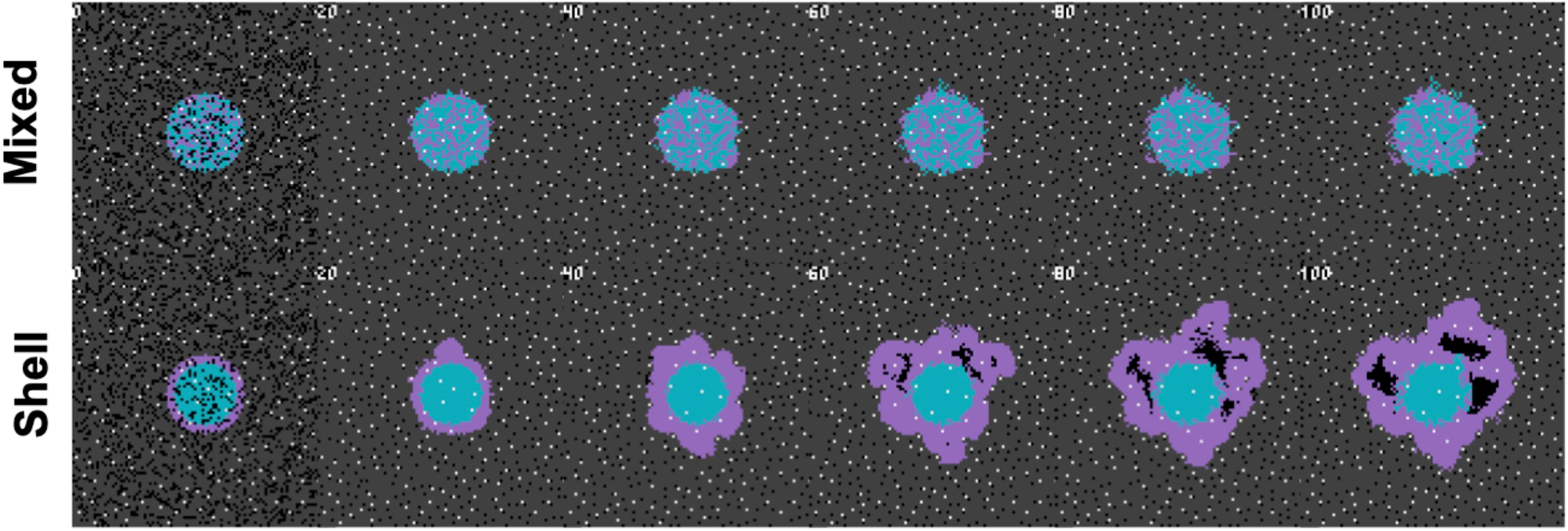
Collective phenotypes drive acid-mediated invasion. Spatial and temporal evolution of two distinct initial spatial configurations of identical numbers of cellular phenotypes leads to differential outcomes due to context-dependent selection. A low glycolysis phenotype (blue) and a high glycolysis phenotype (purple) compete for resources according to the rules outlined in section S5. Top row: a mixed configuration leads to no evolution. Acid-mediated invasion does not occur because the volumetric concentration of acid produced by aggressive cells is not enough to cause invasion when highly glycolytic cells are seeded far apart. Bottom row: In contrast, artificially placing the aggressive high glycolysis phenotypes on the rim leads to invasion from increased volumetric acid via a group-effect.

### 1.4 Mathematical modeling of immune metaphenotypes

Below, we propose and define six metaphenotypes in the context of immune escape and immunotherapy (see figure 2). Then, we present a hybrid multiscale agent-based mathematical model that incorporates phenotypic, vascular, microenvironmental, and spatial heterogeneity to investigate the evolution of aerobic glycolysis in response to immune predation, over a biologically-realistic temporal scale (figure 3). Next, we model immune predation by T-cells in the metabolically altered tumor microenvironment, including immune escape mechanisms such as acid-mediated inactivation of T-cell activation, T-cell inhibition by checkpoint ligand expression on tumor cells, and T-cell glucose deprivation (figure 4). Finally, we quantify the evolution of metaphenotypes over time, illustrating the explanatory power of collective phenotypes in describing the response to buffer therapy and anti-PD-L1 in mono- and combination therapy (figure 5).

**Figure 2.**
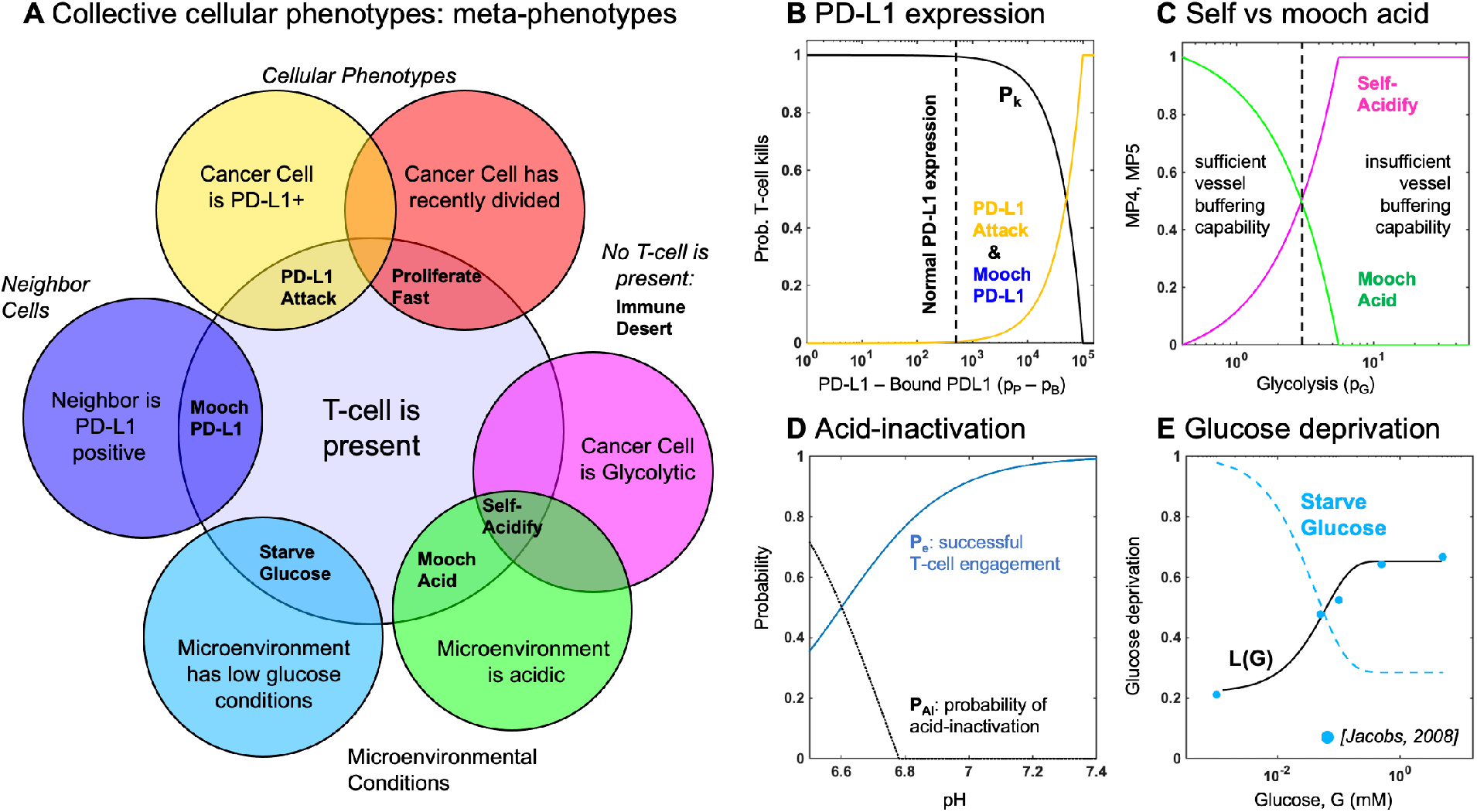
Defining metaphenotypes in the context of immune escape. (A) Six collective cellular metaphenotypes are defined as cancer cells with a given phenotype (e.g. PD-L1), microenvironmental condition (e.g. high acid or low glucose), or neighboring cell. Immune desert is the absence of recent immune interaction. (B) PD-L1 metaphenotypes depend on the likelihood of T-cell kill as a function of PD-L1 expression of self (**PD-L1 Attack**) or neighbor (**Mooch PD-L1**). (C) Acidification metaphenotypes depend on the rate of acidification contributed by self (**Self-Acidify**) or neighbors (**Mooch acid**). (D) The rate of acid-inactivation of T-cells. (E) Data from ref. 34 (blue dots) was used to parameterize T-cell death rate in low glucose, shown in eqn. 20. The **Starve Glucose** metaphenotype expression corresponds to low glucose concentrations.

**Figure 3.**
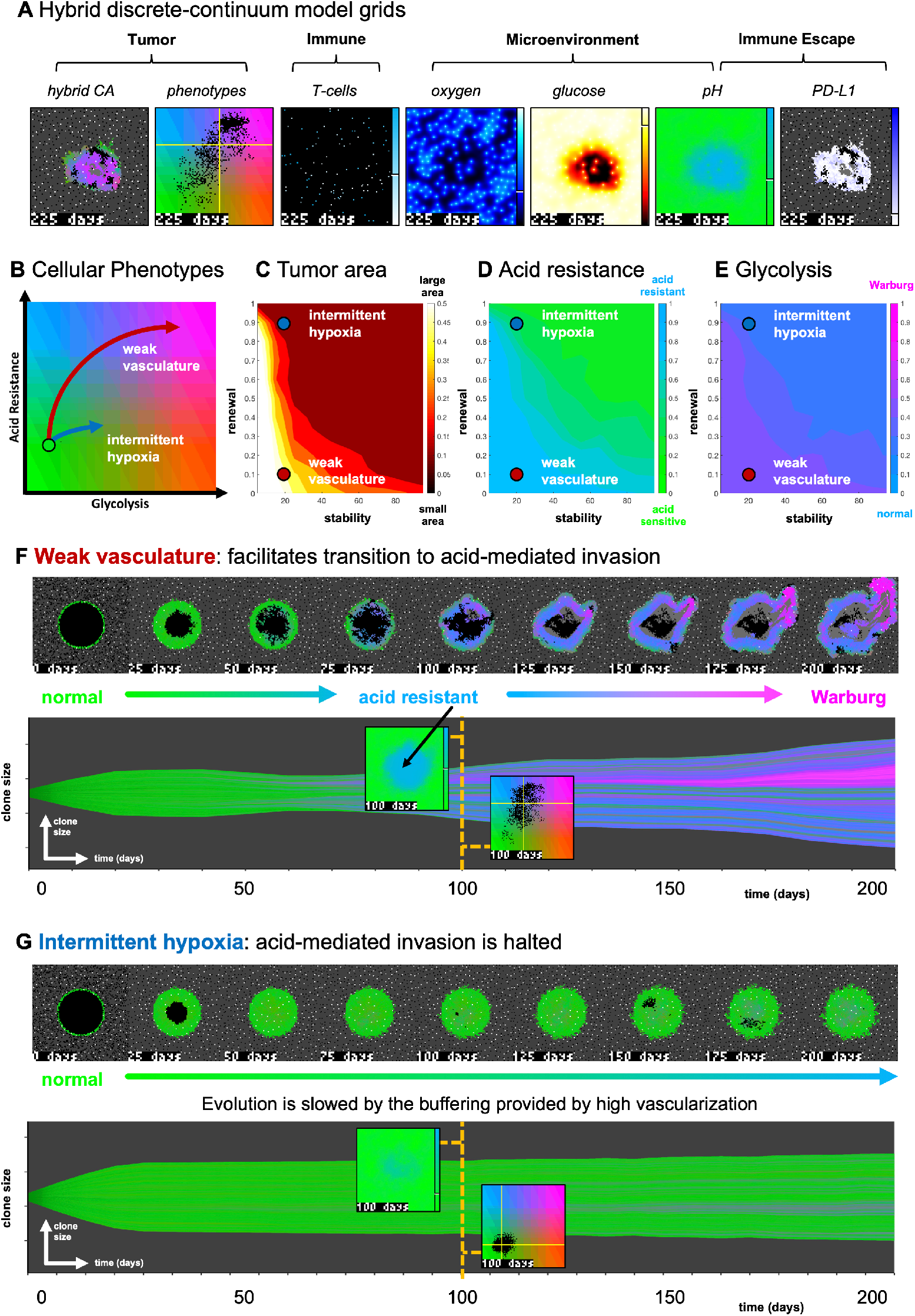
The effect of vasculature renewal and stability on tumor size and phenotype. (A) An example realization of “weak vasculature” (ν_*mean*_ = 20; *p*_*ang*_ = 0.1). Acidic conditions in tumor core select for acid resistant and glycolytic Warburg phenotype. (B) An example realization of “intermittent hypoxia” (ν_*mean*_ = 20; *p*_*ang*_ = 0.9), where selection is limited because of adequate vascularization within the tumor core. (C,D,E) *N* = 10 stochastic realizations are simulated, and the average tumor area (C), acid resistance phenotype (D), and glycolytic phenotype (E) across 10 values of stability (ν_*mean*_ *∈* [0, 100] days), and 10 values of renewal (*p*_*ang*_ *∈* [0, 1]). See associated supplemental video S2.

**Figure 4.**
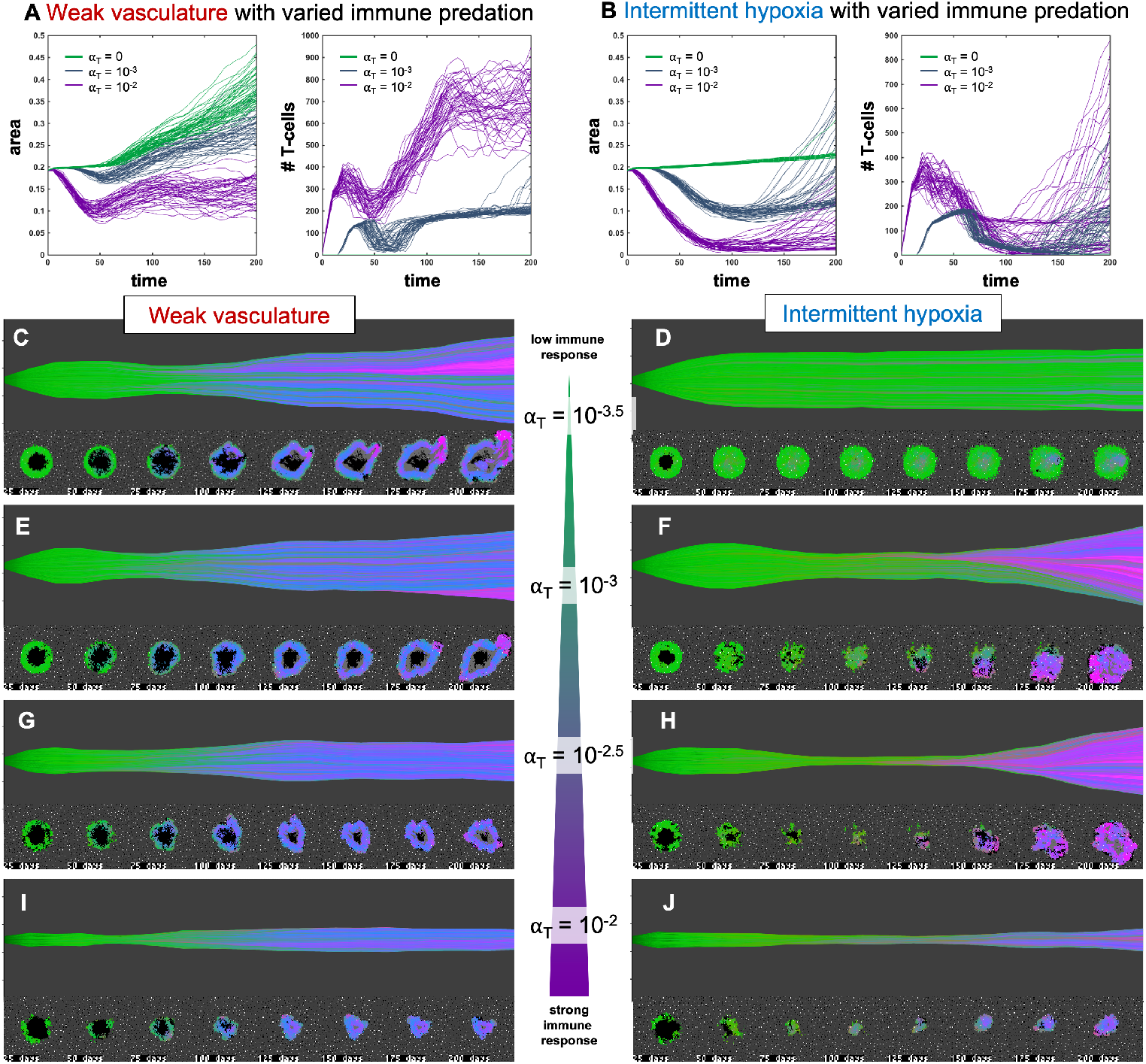
Immune predation induces an evolutionary bottleneck. (A,B) Tumor area over time (left) and the number of T-cells for weak vasculature (A) and intermittent hypoxia (B) conditions), shown for no T-cells (green; α_*T*_ = 0), medium (blue-gray; α_*T*_ = 10^*−*3^) and high (purple; α_*T*_ = 10^*−*2^) immune response rates. (C-J) Example simulation stochastic realizations are shown across a range of immune response from low (top) to high (bottom). Immune predation tends to suppress tumor growth in weak vasculature conditions. In contrast, immune predation induces an evolutionary bottleneck for medium immune response rates (e.g. see F, H), causing aggressive tumor growth compared to the baseline of no immune response.

**Figure 5.**
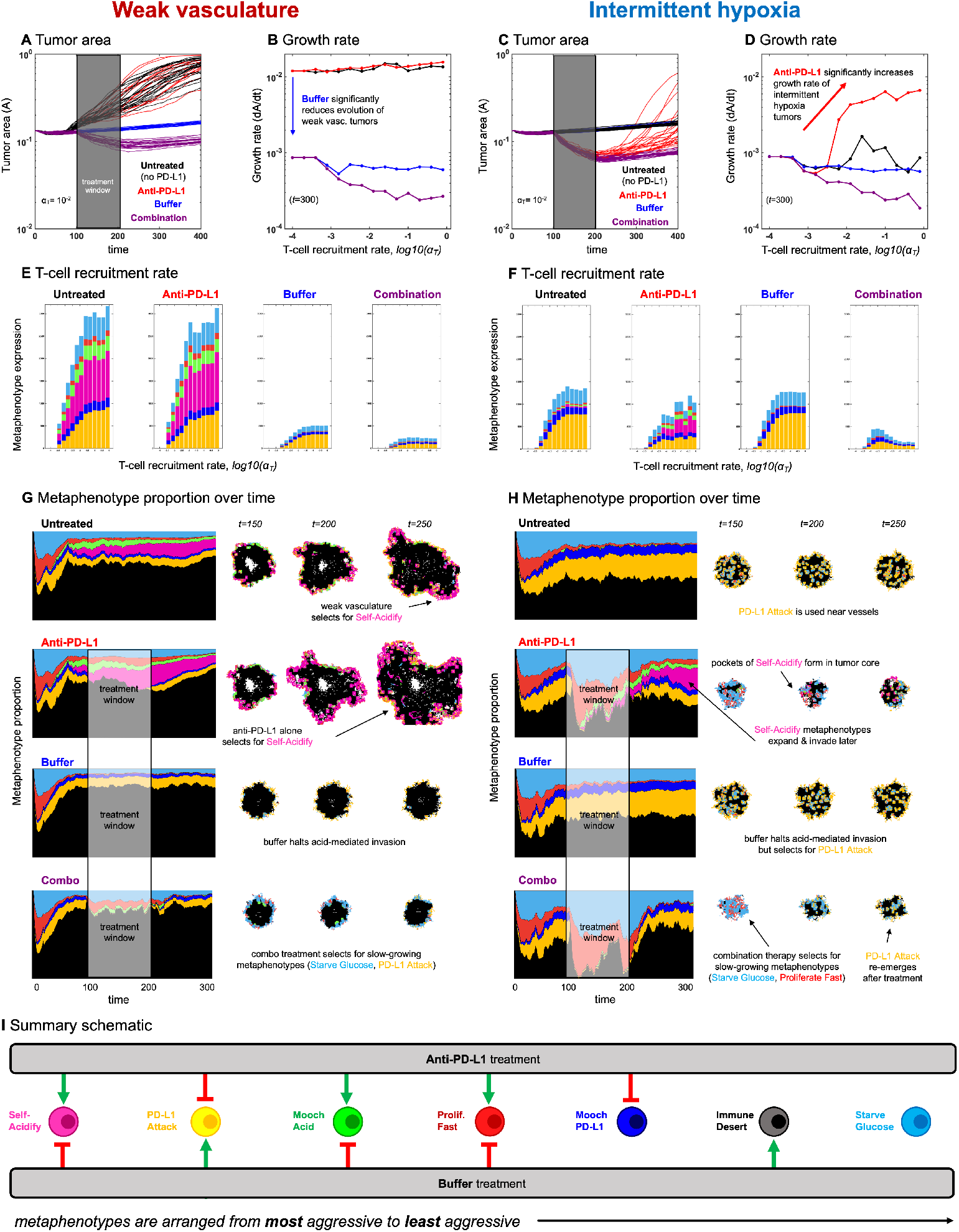
Evolution of metaphenotypes under treatment. Outcomes of tumor response and immune escape can be explained by observing the evolution of metaphenotypes under treatment with anti-PD-L1 (red) and buffer (blue), given in isolation or combination (purple). (A) Tumor area over time (weak vasculature) (B) growth rate over time (weak vasculature). (C) Tumor area over time (intermittent hypoxia vasculature) (D) growth rate over time (intermittent hypoxia vasculature). (E,F) Final distribution of metaphenotypes (exlcuding **Immune Desert**, see fig. S6) at *t* = 300, repeated for weak vasculature (E) and intermittent hypoxia (F). (G,H) Muller plots showing the frequency of metaphenotypes over time in untreated and mono-or combination therapy, with snapshots of spatial configurations during and after treatment, with moderate immune predation (α_*T*_ = 10^*−*2^). See associated supplemental videos S3 and S4 and figure S6.

## 2 Methods

### 2.1 Defining collective cellular phenotypes: immune metaphenotypes

First, we define six collective phenotypes (metaphenotypes) defined as a mechanism of immune escape (see Venn diagram in figure 2A). Each metaphenotype is contingent on a recent tumor-immune interaction and defined in the context of local microenvironment, with the exception of a “null” metaphenotype: **Immune Desert**. The “null” metaphenotype is the lack of collective behavior: **Immune Desert** are cells which do not interact with T-cells. Next, we quantify two PD-L1 metaphenotypes: a counter-attack (tumor cell with high PD-L1 expression that has recently interacted with a T-cell; **PD-L1 Attack**, yellow), and a mooching PD-L1 (**Mooch PD-L1**, blue). As seen in figure 2B, **PD-L1 Attack** is high in cells with high PD-L1 expression while **Mooch PD-L1** is high in cells with low PD-L1 expression, but with neighbors that are high in **PD-L1 Attack**. See Box 1, equations 3 and 4. Two metaphenotypes rely on acid-inactivation: self-acidifying (highly glycolytic cells which secrete acid; **Self-Acidify**, pink) and non-producers (reside in acidic niche but do not produce acid; **Mooch Acid**, green). As seen in figure 2C, **Self-Acidify** is high in cells with a high glycolytic phenotype, hence high acid production (see 5). In contrast, **Mooch Acid** cells have low glycolytic phenotype (not producing acid) but reside in highly a acidic niche that inactivates T-cells (figure 2D). See Box 1, equations 6 and 7. We also consider a proliferative phenotype that has recently divided into empty space (**Proliferate Fast**; red). See Box 1, equations 8. Tumor cells also compete with immune cells for glucose (**Starve Glucose**; light blue). Figure 2E illustrates that **Starve Glucose** reside in areas with a high probability that T-cells die due to low glucose concentration. See Box 1, equations 9. Importantly, each of these metaphenotypes (excluding **Immune Desert**) is contingent on a recent tumor-immune interaction, allowing us to track *effective* collective phenotypes: only metaphenotypes which survive an immune interaction.

#### Box 1

Defining metaphenotypes

Let *T* (*x, y*) be a two-dimensional grid representing the time since the last T-cell interaction has occurred within the local neighborhood of grid location (*x, y*). We define the tumor-immune interaction grid, *I*(*x, y*), where *I* = 1 if an immune cell has traversed within a cancer cell’s neighborhood within the previous *T*_*w*_ days and *I* = 0 otherwise at the current timestep, *t*.

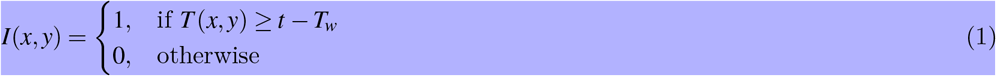

Metaphenotypes (MP) are defined in such a way that MP expression is scaled from zero to one and each cell can take on multiple MP: *M-* = {*m*_1_, *m*_2_,…, *m*_7_} where *m*_*i*_ ∈ [0, 1]

##### 2.1.1 MP1: Immune Desert

We first consider the absence of immune interaction: the immune desert metaphenotype, MP1:

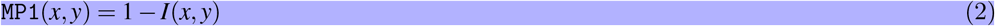

##### 2.1.2 MP2: PD-L1 Attack

Next, we classify cells which employ the PD-L1 counter-attack, defined as high PD-L1 expression (low probability of T-cell kill; see equation 17) with a recent T-cell interaction:

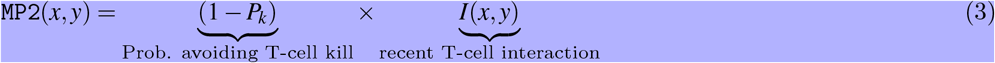

##### 2.1.3 MP3: Mooch PD-L1

In contrast to MP2, cells which interact with T-cells but have low PD-L1 expression can rely on (“mooch”) neighboring cell protection. Here, the metaphenotype is proportional to neighborhood PD-L1 expression.

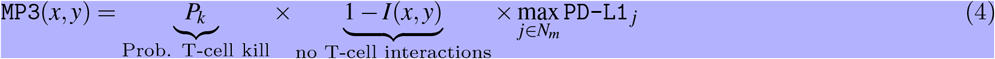

where *N*_*m*_ is a Moore neighborhood of *N*_*m*_ = 8 neighbors.

##### 2.1.4 MP4: Self-Acidify

As cell increase glycolytic capacity (phenotype value *p*_*G*_), more protons are added. The per cell proton production rate is given by:

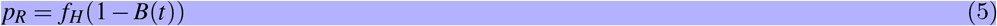

where proton production (see Methods eqn. 14) is scaled by buffer treatment concentration, *B*(*t*).

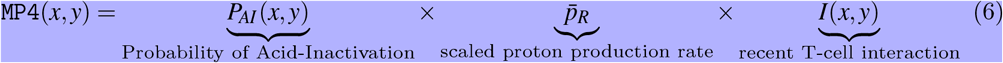

where the production rate, 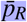, is normalized such that any value for phenotype above the buffering capability of a vessel is assumed to be mostly self-acidify metaphenotype (MP4), while below is assumed to be mostly mooch acid (MP5).

##### 2.1.5 MP5: Mooch Acid

Similarly, the mooch acidify metaphenotype occurs when the probability of T-cell acid-inactivation is high, but where the highly acidic microenvironment is not due to self-acidification.

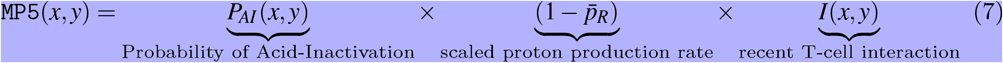

This metaphenotype typically occurs early in simulations in empty regions without tumor or vasculature.

##### 2.1.6 MP6: Proliferate Fast

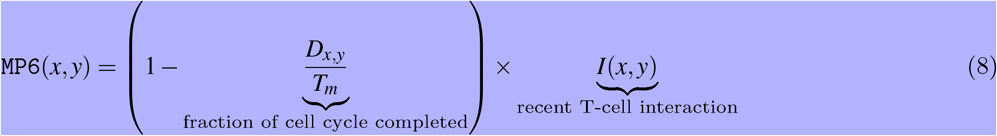

where *D*_*i*_ is the time until next division for the cell at location (*x, y*) and *T*_*m*_ is the inter-mitotic cell division time for a metabolically normal cell.

##### 2.1.7 MP7: Starve Glucose

Tumor cells may also compete with T-cells to starve immune cells of glucose, giving rise to the following metaphenotype:

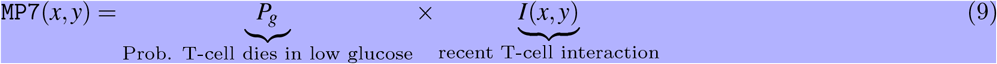

#### 2.2 Hybrid discrete-continuum multiscale model

We test the validity of these metaphenotypes in explaining tumor-immune interactions using a hybrid discrete-continuum multiscale model built using the Hybrid Automata Library framework^32^. The mathematical model here is an extension of an experimentally validated multiscale model of cancer metabolism that incorporates the production of acid and acquired resistance to extracellular pH^5–7, 33^. Figure 3A visualizes the model, which simulates a two-dimensional slice (panel A) through a tumor via a coupled cellular automata and partial differential equation model. A snapshot of multi-scale hybrid cellular automata model is shown (left-to-right) of the tumor spatial map, phenotypes, T-cells, diffusible molecules (oxygen, glucose, acid), PD-L1 and immune susceptibility. Values for parameterization are shown in Supplemental Table 1. Values for parameters are typically identical to previous publications using the non-immune metabolism model^5, 6^, except where parameter values are shown in brackets. In these cases, a parameter sweep is performed across the full range in order to determine the effect of the parameter on outcomes and test hypotheses. For convenience, we re-write the full model description, rules, and cell behaviors in section S5.

## 3 Results

### 3.1 The effect of vasculature renewal and stability on tumor size and phenotype

In figure 3, simulations are shown with the absence of immune predation to establish the model’s baseline dynamics, before quantifying immune predation in the next figure. The model tracks two tumor phenotypes: acid resistance and glycolysis (figure 3B), which vary according to vascularization settings. We compare two classifications of vasculature: weak vasculature (associated with low vessel stability and low rates of vessel renewal) and intermittent hypoxia (associated with low stability, but high renewal). Panels C, D, and E show the average tumor area (C), and tumor phenotypes (D,E) for simulations across a range of vascular settings (no immune). Weak vasculature (low stability and renewal) typically results in larger tumors, more acid resistant phenotypes, and highly glycolytic phenotypes. Weak vasculature induces an acidic niche in the tumor core, selecting for acid-resistant phenotypes (blue). Increased turnover enables increased evolution and selection for aggressive Warburg phenotypes (pink), leading to acid-mediated invasion into surrounding normal tissue. Intermittent hypoxia (low vascular stability with high rates of renewal) generally leads to lower rates of selection and subsequently less invasion (figure 3B).

Spatial maps of phenotypes are shown over time in figure 3F, G along with a visualization called “phenotypic barcoding”, which visualizes the clone size, phenotype and lineage information over time^7^ using the EvoFreq R package^35^ (for more information on interpreting phenotypic barcoding plots, see figure S5). Figure 3A depicts the process by which weak vasculature selects for aggressive tumor growth. Acidic conditions in the tumor core (low glucose, low oxygen, and high pH) cause rapid death of glycolytically normal tumor cells with low levels of acid resistance. Selection for acid resistance occurs first (blue phenotypes), followed by selection for highly glycolytic tumor cells (pink phenotypes) which eventually invade into surrounding tissue. Conversely, in figure 3B, intermittent hypoxia conditions result in little selection. The well-vascularized tumor core limits selection for aggressive phenotypes. This result underscores the importance of understanding the baseline vascular conditions before modeling the complex dynamics with the additional immune predation. A snapshot of the intratumoral oxygen, immune susceptibility (see equation 21), phenotypes, and pH is shown at the end of each simulation.

### 3.2 Immune predation induces an evolutionary bottleneck

Figure 4 shows the response of two vascular conditions (weak and intermittent hypoxia) under no immune response (green; α_*T*_ = 0), medium (blue-gray; α_*T*_ = 10^*−*3^) and high (purple; *α*_*T*_ = 10^*−*2^) immune response rates. Immune cells are recruited in proportion to tumor size and response rate, *α*_*T*_.

Immune response tends to suppress tumor growth in weak vasculature conditions (figure 4A, left). Compared to baseline tumor growth, all levels of immune response result in greater tumor suppression. In contrast, immune predation in intermittent hypoxia conditions often leads to an initial response but fast regrowth (figure 4B, left). This is confirmed by visual inspection of the phenotypic barcoding visualizations in figure 4C-J. Weak vascular conditions select for aggressive phenotypes with little-to-no immune presence (figure 4C). The addition of immune cells only serves to slow an already aggressive tumor (figure 4E,G,I). In stark contrast, intermittent hypoxia conditions rarely select for strong growth in the absence of immune predation (figure 4D). Immune predation serves as a selection pressure, in conditions where there would otherwise be very little selection.

Immune predation under intermittent hypoxia conditions induces an evolutionary bottleneck for medium immune response rates (e.g. see F, H), causing fast selection for aggressive growth compared to the baseline of no immune response. Interestingly, this effect occurs on a “Goldilocks” scale. The long neck of the bottleneck is associated with higher rates of tumor turnover (due to immune attack), selecting for phenotypes which are 1) inside an immune-evasive niche or 2) rapidly divide to outgrow immune kill.

### 3.3 Metaphenotypes explain immune escape under treatment

After establishing the baseline dynamics without (fig. 3) and with (fig. 4) immune predation, we next consider two treatments to mitigate immune escape and to reduce tumor growth: anti-PD-L1 and a pH buffer given in isolation or combination. A short window of treatment is simulated and results are compared to the untreated baseline. As seen in figure 5A-D, combination therapy outperforms monotherapy in both vascular settings, but vascular dynamics drive differences in monotherapy outcomes. For example, anti-PD-L1 (red) therapy does not appreciably slow tumor evolution or growth in weak vasculature (fig. 5A,B). In contrast, anti-PD-L1 does induce large tumor remission in intermittent hypoxia (fig. 5C,D), albeit only temporarily before a strong relapse. These results are seen across a range of immune recruitment rates (fig. 5B,D).

The metaphenotypes leading to immune escape are shown in figure 5E,F for each treatment scenario. As T-cell recruitment rate increases left-to-right, tumors evolve metaphenotypes in response to immune infiltration. Vascularization drives differential selection of metaphenotypes in baseline untreated dynamics. Weak vasculature (panel E; untreated) is associated with acidification metaphenotypes (**Self-Acidify**, pink; **Mooch Acid**, green). These are aggressive, highly glycolytic metaphenotypes that facilitate acid-mediated invasion. In contrast, intermittent hypoxia (panel F; untreated) selects for PD-L1-based immune-escape mechanisms (**PD-L1 Attack**, yellow; **Mooch PD-L1**, dark blue).

Treatment alters the type and magnitude of metaphenotype expression. Anti-PD-L1 selects for acidification metaphenotypes (self-acidify or mooch acid) in both vascularization cases. Buffer treatment eliminates the emergence of both **Self-Acidify** and **Mooch Acid** phenotypes by slowing evolution (e.g. refer to fig. 3B). But in response, **PD-L1 Attack** is selected (yellow). Only combination therapy targets both acidification metaphenotypes and PD-L1 phenotypes. Tracking the response of metaphenotypes to treatment explains why combination therapy is ideal for minimizing tumor growth, compared to monotherapy options. Importantly, only combination decreases the sum total of metaphenotypes expressed, and specifically targets aggressive phenotypes (**Self-Acidify** and **Mooch Acid**) across both vascularization scenarios.

### 3.4 Spatial configuration of metaphenotypes under treatment

The explanatory power of these defined metaphenotypes is seen most clearly by observing their spatial arrangement under high immune predation (see figure 5G,H and associated Supplemental Videos S2 and S3). For example, weak vasculature (fig. 5G) is associated with the **Self-Acidify** and **PD-L1 Attack** metaphenotypes on the invasive front of the tumor. Much of the tumor interior is unaffected by immune cells (**Immune Desert**), regardless of tumor phenotype. Treatment with Anti-PD-L1 selects for the aggressive **Self-Acidify** metaphenotype, while Buffer selects for **PD-L1 Attack** on the tumor rim. Combination therapy is required to achieve maximum tumor response, resulting in small tumors consisting mostly of non-aggressive metaphenotypes (**Starve Glucose** or **Proliferate Fast**).

In contrast, under intermittent hypoxia vasculature the **Immune Desert** comprises a much lower fraction of tumor metaphenotypes, as this improved vascularization delivers T-cells into the tumor core. **PD-L1 Attack** is used near blood vessels and on the tumor rim, and **Self-Acidify** does not occur due to low turnover in untreated conditions. Treatment with Anti-PD-L1 negates immune escape from **PD-L1 Attack**, inducing cellular turnover and subsequently selecting for **Self-Acidify** and **Mooch Acid** metaphenotypes. Combination therapy results in small, slow-growing tumors with less aggressive metaphenotypes (**Mooch PD-L1** and **Starve Glucose**).

In both vasculature settings, cells slightly inset from the rim use metaphenotypes that **Mooch Acid** and **Mooch PD-L1** from cells on the rim (see Supplemental Videos S2 and S3) while cells in regions of high turnover employ the **Proliferate Fast** metaphenotype. **Starve Glucose** remains at low levels throughout all treatment modalities and vasculature settings. As seen in the supplemental videos (S2 and S3), it is difficult to determine the major driver of immune escape from the maps of phenotypes alone, as areas of high glycolysis and high PD-L1 are each spatially heterogeneous and overlapping.

## 4 Discussion

The importance of acidity in modulating immune response in cancer is only just beginning to be understood. Our results highlight the potential utility in buffering agents combined with immunotherapy. Whilst such agents are not currently used in cancer treatment there is a growing body of evidence that more alkaline diets can facilitate standard cancer treatments. Patient compliance with sodium bicarbonate has been an an issue in previous clinical trials due to GI irritability, leading to an investigation of dietary intake of highly buffered foods or supplements^36^. The exact mechanism of action is as yet to be understood but previous work has shown how acid-mediated invasion can be modulated through diet changes^36^. Several factors contribute to a lack of responsiveness to immune checkpoint blockers, including abnormal tumor microenvironment where poor tumor perfusion hinders drug delivery and increases immunosuppression^37^. Poor vascularization also leads to a hypoxic and therefore acidic microenvironment, increasing immunosuppression. The modeling above recapitulates this trend, as immune predation is less effective in weak vascularized tumors than in intermittently vascularized tumors. Vascular renormalization can be enhanced through administration of anti-angiogenic agents (e.g., anti-vascular endothelial growth factor agents) to fortify immature blood vessels and improve tumor perfusion^38^. However, our results indicate that administration of immune checkpoint blockade in tumors with increased vascularization may lead to a short-term good response but poor long-term outcomes as selection for increased glycolysis occurs. Mathematical modeling allows for direct comparison of initially identical simulations in the absence (fig. 3) and presence (fig. 4) of immune predation. We observe an immune gambit under high vascular renewal (intermittent hypoxia), due to an evolutionary bottleneck. The impact of this evolutionary bottleneck is reduced when anti-PD-L1 is combined with buffer therapy.

Characterization of collective phenotypes into metaphenotypes enables a straightforward explanation of the effect of treatment in a complex, multi-scale model. This characterization is necessary, in part, due to the fact that acid-mediated invasion is a collective phenotype phenomenon (fig. 1). Immune escape is also, by definition, a collective phenomenon by requiring the presence of two cell types in close proximity: tumor and immune. A summary schematic of the results is shown in figure **??**. The interaction diagram describes the role of anti-PD-L1 and buffer in either promoting (green) or inhibiting (red) each metaphenotype. Broadly, the two treatments offset one another by inhibiting the metaphenotypes that the opposite treatment promotes. The two exceptions, starve glucose and immune desert, are both non-aggressive phenotypes. This summary schematic illustrates the utility of defining metaphenotypes in the context of treatment to provide insight into immune-escape dynamics. The most dominant mechanism of immune escape seen in the model is the lack of immune interactions (immune desert), especially when the tumor bed is poorly vascularized. Tumor-expressed PD-L1 is a viable immune-escape mechanism in the absence of treatment, across a range of vascularization, but treatment with anti-PD-L1 selects for the two acid-inactivation metaphenotypes (**Self-Acidify** and **Mooch Acid**). Environmental conditions must also consider neighboring (and self) cellular phenotypes. A cell in acidic conditions may rely on acid-inactivation either by self-production of acid or mooching from neighboring producer cells, a form of “public good”^39^. Buffer therapy limits selection for self-acidification, driving selection toward less aggressive metaphenotypes (**Glucose Starvation** or **Immune Desert**). It’s also important to note that mooching metaphenotypes only occur in the presence of non-mooching phenotypes. Because of this, and the fact that phenotypes of individual cells change only slowly (upon division), mooching phenotypes are not expected to be a viable long-term immune escape strategy, but limited to transient, local patches co-localized with non-moochers. However, in a model where the ratio of two phenotypes is determined stochastically, for example, a population of both phenotypes could coexist for a longer period of time.

The intimate feedback between a growing tumour and the homeostatic tissue its invading drives both ecological and evolutionary dynamics that should not be ignored in modern cancer therapy. The results we presented here indicate that treatments that modulate context may turn out to be just as important as those that target the tumour.

## Acknowledgments

The authors gratefully acknowledge funding by the National Cancer Institute via the Cancer Systems Biology Consortium (CSBC) U01CA232382, the Physical Sciences Oncology Network (PSON) U54CA193489, and support from the Moffitt Center of Excellence for Evolutionary Therapy.

## Author Contributions

ARAA and MRT conceived the research questions and model design. KAL contributed to model design. JW, FR, RB, and CA performed the modeling and analyses. All authors contributed to writing and editing the manuscript.

