## Supplemental Information for "Tumor-immune metaphenotypes orchestrate an evolutionary bottleneck that promotes metabolic transformation"

### S5 Hybrid discrete-continuum multiscale model

The mathematical model here is an extension of a published multi-scale model of cancer metabolism[5–7,33]. Figure S1A visualizes the model, which simulates a two-dimensional slice (panel A) through a tumor via a coupled cellular automata and partial differential equation model. Snapshots (left-to-right) of the tumor spatial map, phenotypes, vascular renewal probabilities, T-cells, diffusible molecules (oxygen, glucose, acid), PD-L1 and immune susceptibility. The model prescribes the behavior of cells interacting with neighbors and environmental factors (figure S1B), the rules governing internal tumor cell decisions (figure S1C), the range of phenotype space (acid resistance, glycolysis, and PD-L1 in figure S1D), and the rules governing T-cell internal decision (figure S1E). Parameterization is shown below in Supplemental Table 1, where chosen parameters are typically identical to previous publications using the non-immune metabolism model[5,6], except where parameter values are shown in brackets. A sweep is performed across the full range of the bracketed parameters in order to determine the effect of the parameter on outcomes and test hypotheses. New parameters developed in this manuscript (i.e. the immune module) are shown below the solid line.

### Box 2: Mathematical model description

#### S5.1 Baseline mathematical model

An overview of the mathematical model is shown in figure S1. The model simulates a two-dimensional slice through a tumor via a coupled cellular automata (CA) and partial differential equation (PDE) model. Vasculature is modeled as a set of point sources, with spacing consistent with those measured in normal stroma[5]. As timescales of metabolism and cell-scale dynamics (e.g. proliferation) vary significantly, we solve the PDEs to reach steady state between CA timesteps. The model includes several diffusible molecules: oxygen, acid, and glucose (figure S1B, yellow boxes). The concentration of each diffusible is modeled by the following diffusion-production-consumption equation:

$$\frac{\partial C}{\partial t} = D\nabla^2 C + f(x,t) \tag{10}$$

where $C$ represents the diffusible molecule concentration, $D$ is the diffusion constant, and $f(x,t)$ is the molecule-specific rate of consumption/production of each particular molecule. For example, oxygen consumption ($f_O$) by cells is given by Michaelis-Menten dynamics:

$$f_O = -V_O \frac{O}{O + k_O} \tag{11}$$

where $k_O$ is the oxygen concentration for half-maximal oxygen consumption and $V_O$ is the maximal oxygen consumption by cells. Glucose consumption is given by the following modified Michaelis-Menten equation:

$$f_G = -\left(\frac{p_G A_o}{2} + \frac{27 f_O}{10}\right)\frac{G}{G + k_G} \tag{12}$$

where $p_G$ is the heritable trait which represents aerobic glycolysis and its resultant excessive glucose consumption. $A_o$ is the ATP production rate in normal cells, $k_G$ is glucose concentration for half-maximal glucose consumption. ATP production rate is given by:

$$f_A = -\left(2 f_G + \frac{27 f_O}{5}\right), \tag{13}$$

and proton production rate given by:

$$f_H = k_H\left(\frac{29(p_G V_O + f_O)}{5}\right). \tag{14}$$

#### S5.2 Phenotypic drift

Both normal and tumor cells undergo a decision process in figure S1C, involving cell cell cycle dynamics, proliferation and phenotypic drift. Cells with low ATP efficiency or cells which are maladapted for acidosis die. Cells take on three phenotypic traits: acid resistance ($p_H$), glycolysis ($p_G$), PD-L1 expression ($p_P$). Phenotypes may drift upon cell division as follows:

$$p_i(t + \Delta t) = p_i(t) \cdot (1 + \Delta_i)^{(U(-1,1))} \tag{15}$$

where $i \in \{H, G, P\}$, representing acid-resistance, glycolysis, PD-L1 phenotypes respectively (see Table 1). $U(-1,1)$ represents a uniform probability distribution drawn on the interval -1 to 1 and $\Delta t$ is the timestep (2 hours). Phenotype variation rate parameters ($\Delta_i \in [\Delta_G, \Delta_H, \Delta_P]$) are shown in Table 1.

#### S5.3 Immune recruitment model

Immune cells are recruited in proportion to the current tumor size a few days prior, $N(t-\tau_T))$ at a rate $\alpha_T$ until the number of T-cells equals or exceeds this value. T-cells undergo a natural decay rate if they have not encountered a tumor cell in the past $\beta_T$ number of days.

$$T(t+1) = \begin{cases} \alpha_T N(t-\tau_T) - \dfrac{1}{\beta_T} T(t), & N(t-\tau_T) > T(t) \\ T(t) - \dfrac{1}{\beta_T} T(t), & N(t-\tau_T) \leq T(t) \end{cases} \tag{16}$$

Tumor cells have several mechanisms for immune evasion in the mathematical model: PD-L1 and acid inactivation.

### S5.4 PD-L1

The probability, $P_k$, that a tumor cell is successfully eliminated by a T-cell is a function of the constitutive PD-L1 expression ($p_P$) and the bound PD-L1 ($p_B$; see section on treatment with anti-PD-L1 below):

$$P_k = 1 - \frac{10^{p_P - p_B}}{10^{p_{P,norm} - p_{P,min}}} \tag{17}$$

where $p_{P,min}$ and $p_{P,norm}$ are the maximal PD-L1 phenotype (corresponding to zero probability of kill; see figure 2B) and normal PD-L1 phenotype, respectively. See table 1 for parameter values.

#### S5.5 T-cell viability in high acid

Recent results from Pilon-Thomas et. al. have suggested that acid does not affect T-cell viability but instead impairs activation[23]. Low pH arrests T-cell cytokine and chemokine production (a measure of activation) . Thus, we model probability of successful engagement of a cancer cell by a T-cell depends on the microenvironmental pH:

$$P_e = \frac{1}{1 + e^{-\sigma_p (H - H_p)}} \tag{18}$$

where $H$ is the pH value, $H_p$ is the half-max engagement probability and $\sigma_p$ represents the pH at which engagement probability is half its maximum value. Secondly, exocytosis of lytic granules is impaired in low pH[40], causing increased time to kill targets in low pH. The probability that a tumor cell successfully inactivates a T-cell due to low acid is given by:

$$P_{AI} = 1 - \frac{P_e \Delta t}{d_e \left(1 + e^{-\sigma_e (H - H_e)}\right)} \tag{19}$$

where $d_e$ represents the minimum engagement time duration, modulated by acid concentration, $H$[41]. $H_e$ is the half-max engagement time and $\sigma_e$ is the steepness parameter, and $\Delta t$ is the length of a single time step (2 hours).

T-cells are also assumed to undergo death at higher rates in low glucose concentrations. This component of the model was parameterized using experimental data from ref. 34, where probability of death after two days is fit to the following equation:

$$L(G) = (L_0 - L_i) e^{G L_g} + L_i, \tag{20}$$

where $L_0$ and $L_i$ T-cell survival rate in low and high glucose concentrations, $G$, with steepness parameter $L_g$ (see figure 2E).

#### S5.6 Immune susceptibility

The total immune susceptibility of a cell is the likelihood of a T-cell kill as a function of PD-L1 expression multiplied by the likelihood that a T-cell is not acid-inactivated:

$$S = P_k(1 - P_{AI}), \tag{21}$$

#### S5.7 Treatment

Two treatments are considered: anti-PD-L1 and buffer therapy. Anti-PD-L1 is modelled as a diffusible molecule (eqn. 10), with tumor cell uptake of bound PD-L1 ($p_B$) at rate $D_A$ and natural decay at rate $\gamma_A$ (see Table 1). Bound PD-L1 in each tumor cell is limited to the range $p_B \in [0, p_P]$, where T-cell kill rate, $P_K$, is a function of the difference between constitutively expressed and bound PD-L1 (see figure 2B). Cell internal bound PD-L1 decays at rate $\gamma_P$. Buffer therapy is modeled as a change in proton buffering coefficient, $k_H$:

$$k_H = 0.00025(1 - B(t)), \tag{22}$$

where $B(t)$ is the time-dependent administration of buffer therapy, and the baseline value of the buffering coefficient is $k_H = 0.00025$ (see Table 1 and ref. 5).

#### S5.8 Local Neighborhoods

The model is carried out on a two-dimensional lattice where each tumor, normal, or immune cell occupies a single lattice location, $(x, y)$. The cell's local neighborhood is a set of lattice locations defined in relation to the focal cell's location, defined as $N_m$ ($N_m = 8$ for a Moore neighborhood). When the focal cell undergoes division a daughter cell is placed in a random neighboring grid point and the parent cell remains on the original lattice point. The cell may undergo apoptosis (death) and is removed from the domain. After each generation cells are shuffled and iterate through in random order in future time steps.

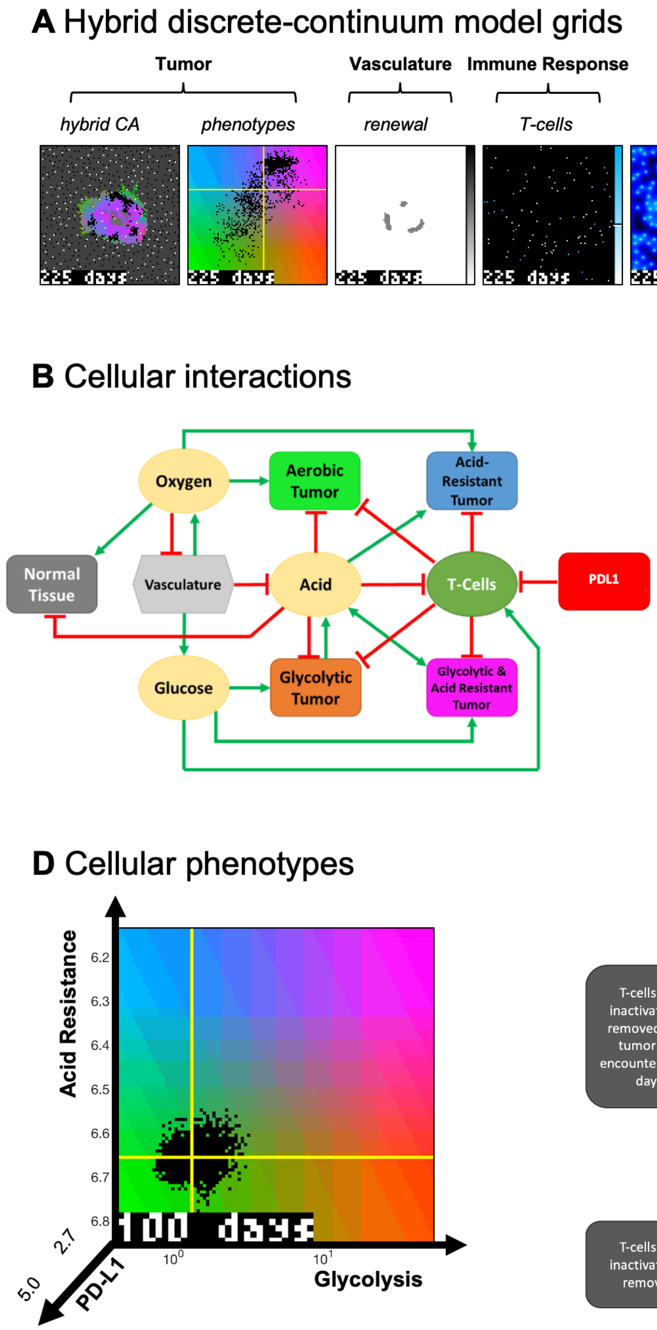


**Figure S1. Hybrid Discrete-Continuum Model Diagram.** (A) Snapshot is shown for example simulation of the full multi-scale hybrid cellular automata model: tumor spatial map, phenotypes, vascular renewal probability, T-cells, diffusible molecules (oxygen, glucose, pH), tumor cell PD-L1, and immune susceptibility (see equation 21). Each snapshot has a corresponding colorbar (right) with a marker indicating average value. (B) Model interaction network for diffusible molecules (yellow), vasculature (light gray), normal tissue (dark gray), and variable tumor phenotypes (colors). Red lines indicate inhibitory interactions while green lines indicated promoting interactions. (C) Decision process for each cell, with diamonds representing decisions. Green arrows indicate that the condition is satisfied, and red that the condition is not met. (D) A map of tumor phenotype state space on three axes. The horizontal axis is the constitutively activated glycolytic capacity ($p_G$), and vertical axis is the change in acid resistance ($p_H$) from normal, with higher resistance to acidic conditions being higher on the plot, and the final axis is constitutively expressed PD-L1 ($p_P$). The normal metabolic phenotype is at the intersection of the two yellow lines, with the cloud of black dots representing normal variation in phenotypes within the tumor composition (each black dot is a single tumor cell). (E) Decision process for T-cells. T-cells are recruited in proportion to tumor size at a rate of $\alpha_T$. T-cells are inactivated and removed if they remain is acidic conditions too long, or if they are inactivated by a PD-L1 positive cancer cell.

**Table 1.** Model Parameterization

| Parameters | Value | Units | Description |
|---|---|---|---|
| $\delta x$ | 20 | $\mu m$ | Diameter of CA grid point |
| $p_D$ | 0.005 | 1/d | Normal tissue death rate |
| $p_\Delta$ | 0.7 | 1/d | Death probability in poor conditions |
| $p_n$ | 5e-4 | 1/d | Necrotic turnover rate |
| $D_O$ | 1820 | $\mu m^2/s$ | Diffusion rate of oxygen |
| $D_g$ | 500 | $\mu m^2/s$ | Diffusion rate of glucose |
| $D_H$ | 1080 | $\mu m^2/s$ | Diffusion of protons |
| $O_O$ | 0.0556 | mmol/L | Oxygen concentration in blood |
| $G_O$ | 5 | mmol/L | Glucose concentration in blood |
| $pH_O$ | 7.4 | pH | pH of blood |
| $V_O$ | 0.012 | mmol/L/s | Maximal oxygen consumption |
| $k_O$ | 0.005 | mmol/L | Half-max oxygen concentration |
| $k_G$ | 0.04 | mmol/L | Half-max glucose concentration |
| $k_H$ | 2.5e-4 | - | Proton buffering coefficient |
| $A_d$ | 0.35 | - | ATP threshold for death |
| $A_q$ | 0.8 | - | ATP threshold for quiescence |
| $p_{H,min}$ | 6.1 | pH | Maximal acid resistance phenotype |
| $p_{H,norm}$ | 6.65 | pH | Normal acid resistance phenotype |
| $\Delta H$ | 0.003 | pH | Phenotype variation rate (acid res.) |
| $p_{G,max}$ | 50 | - | Maximal glycolytic phenotype |
| $\Delta G$ | 0.15 | - | Phenotype variation rate (glycolysis) |
| $\tau_{min}$ | 0.95 | Days | Minimum cell cycle time |
| $\sigma_{min}$ | 80 | $\mu m$ | Minimum vessel spacing |
| $\sigma_{mean}$ | 150 | $\mu m$ | Mean vessel spacing |
| $v_{mean}$ | [5, 100] | Days | Vessel stability |
| $p_{ang}$ | [0, 1] | - | Angiogenesis rate |
| $T_M$ | 1 | - | Probability T-cell moves |
| $\tau_T$ | 4 | - | T-cell response delay |
| $\alpha_T$ | [1e-4,1e-1] | - | T-cell recruitment rate |
| $\beta_T$ | 10 | Days | Non-activated T-cell decay |
| $p_{P,min}$ | 5 | - | Maximal PD-L1 phenotype |
| $p_{P,norm}$ | 2.7 | - | Normal PD-L1 phenotype |
| $\Delta P$ | [0,1] | - | Phenotype variation rate (PD-L1) |
| $d_e$ | 0.042 | Days | T-cell engagement duration |
| $H_e$ | 6.6 | - | half-max pH T-cell engagement time |
| $\sigma_e$ | 4 | - | steepness of T-cell engagement time |
| $H_p$ | 6.6 | - | half-max pH T-cell engagement probability |
| $\sigma_p$ | 6 | - | steepness of T-cell engagement probability |
| $L_i$ | 65.35 | percent | T-cell survival rate in high glucose |
| $L_0$ | 21.78 | percent | T-cell survival rate in low glucose |
| $L_g$ | -16.67 | percent | T-cell glucose deprivation parameter |
| $D_A$ | 100 | $\mu m^2/s$ | Anti-PD-L1 diffusion parameter |
| $\gamma_A$ | 0.5 | $1/s$ | Anti-PD-L1 natural decay rate |
| $\gamma_P$ | 0.001 | $1/s$ | Cellular bound PD-L1 decay rate |

**Figure S2** **Supplemental Video S1.mov** Simulation of the full multi-scale hybrid cellular automata model: tumor spatial map, phenotypes, vascular renewal probability, T-cells, diffusible molecules (oxygen, glucose, pH), tumor cell PD-L1, and immune susceptibility (see equation 21). Each grid has corresponding colorbar (right) with marker indicating average value. This is an example realization of "weak vasculature" ($\nu_{mean} = 20; p_{ang} = 0.1$). Acidic conditions in tumor core select for acid resistant and glycolytic Warburg phenotype (pink color on the Tumor grid). Subsequently, nectrotic cells form in the tumor interior. Immune response is weak ($\alpha_T = 10^{-3}$), not selecting for PD-L1 positivity. The primary means of immune escape is acid-inactivation of T-cells caused by acid-producing Warburg cells.

**Figure S3** **Supplemental Video S2.mov** Evolution of metaphenotypes in **Weak** vasculature under high immune predation ($\alpha_T = 10^{-2}$). The left-hand side shows three phenotypes (Acid Resistance, Glycolsis, and PD-L1), along with acid concentration and T-cell location. It is difficult to determine the major driver of immune escape from the maps of phenotypes alone: areas of high glycolysis and high PD-L1 are each spatially heterogeneous and overlapping. However, metaphenotypes give a detailed picture of immune escape dynamics. Much of the tumor interior is unaffected by immune cells (**Immune Desert**), regardless of tumor phenotype. The outer rim is immune-protected by **PD-L1 Attack** and **Self-Acidify** phenotypes. Slightly inset from the rim, cells use metaphenotypes that **Mooch Acid** and **Mooch PD-L1** from cells on the rim. Cells in regions of high turnover employ the **Proliferate Fast** metaphenotype. **Starve Glucose** remains at low levels throughout all treatment modalities. Treatment with Anti-PD-L1 selects for the aggressive **Self-Acidify** metaphenotype, while Buffer selects for **PD-L1 Attack**. Combination therapy results in small, slow-growing tumors with less aggressive metaphenotypes (**Mooch PD-L1** and **Starve Glucose**).

**Figure S4** **Supplemental Video S3.mov** Evolution of metaphenotypes in **Intermittent Hypoxia** vasculature under high immune predation ($\alpha_T = 10^{-2}$). The left-hand side shows three phenotypes (Acid Resistance, Glycolsis, and PD-L1), along with acid concentration and T-cell location. Similar to S3, it is difficult to determine the major driver of immune escape from the maps of phenotypes alone: areas of high glycolysis and high PD-L1 are each spatially heterogeneous and overlapping, but metaphenotypes give a detailed picture of immune escape dynamics. Here, **Immune Desert** comprises a much lower fraction of tumor metaphenotypes, as better vascularization delivers T-cells into the tumor core. **PD-L1 Attack** is used near blood vessels and on the tumor rim, with **Mooch PD-L1** employed by nearby cells. In untreated conditions, **Self-Acidify** does not occur due to low turnover. However, Anti-PD-L1 negates immune escape from **PD-L1 Attack**, inducing turnover and selecting for **Self-Acidify** and **Mooch Acid** metaphenotypes. Combination therapy results in small, slow-growing tumors with less aggressive metaphenotypes (**Mooch PD-L1** and **Starve Glucose**).

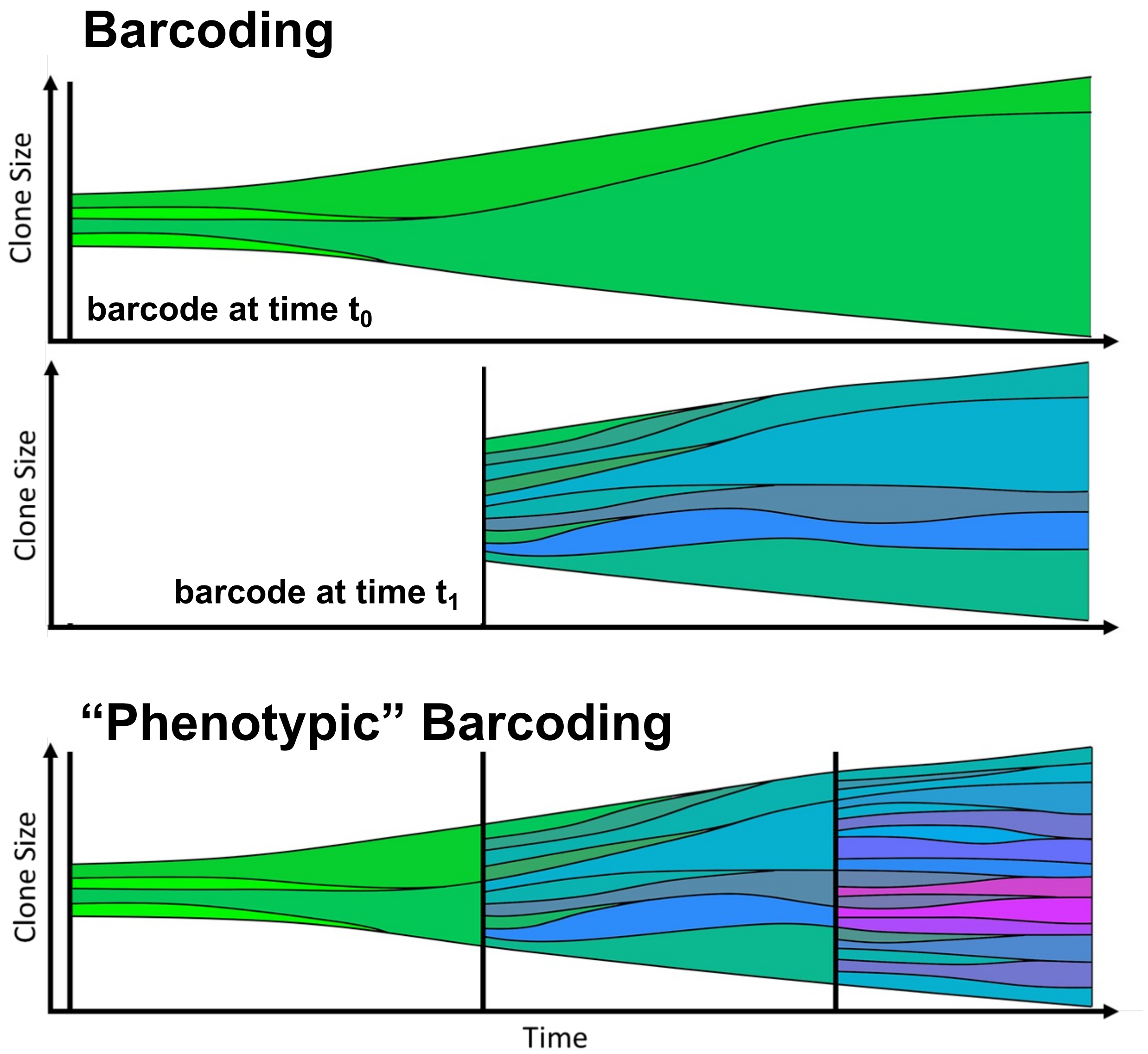


**Figure S5. Phenotypic barcoding** Schematic of “phenotypic barcoding” scheme for visualizing tumor evolution. At time point 0, all cells are given a unique ID, also known as a barcode (top). This can be repeated at later times (e.g., time point t1) by adding a second unique ID to each extant cell (middle). Clones and subclones can be re-colored by average phenotype (bottom) so that both the phenotype and lineage information are visualized during tumor evolution.

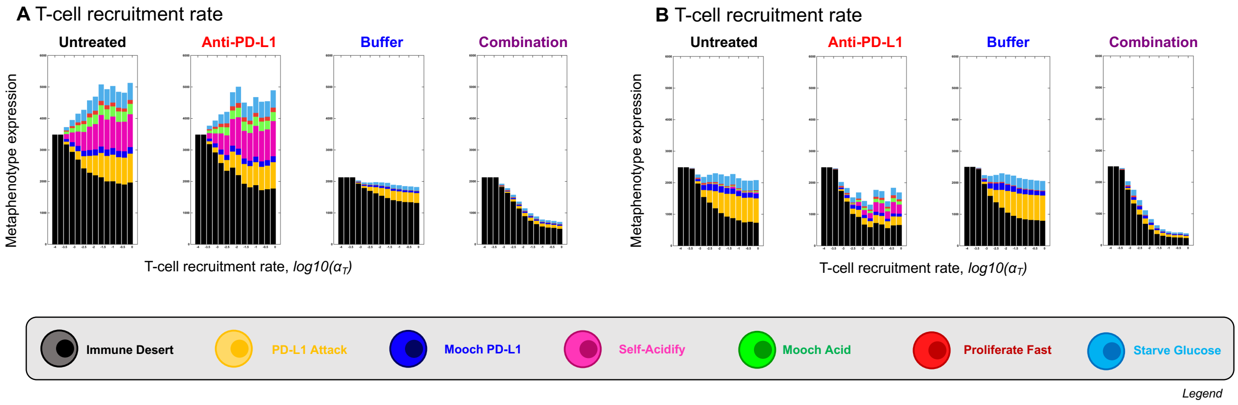


**Figure S6. Evolution of metaphenotypes under treatment.** Supplementary figure corresponding to figure 5E,F in the main text. (A) Final distribution of metaphenotypes after treatment ($t = 300$; weak vasculature), including **Immune Desert**. (B) Final distribution of metaphenotypes after treatment ($t = 300$; intermittent hypoxia), including **Immune Desert**. In all treatment scenarios, the presence of **Immune Desert** phenotypes decreases as T-cell recruitment rate increases (left-to-right), indicative of increased T-cell perfusion.
